# Prospective control of movement in the basal ganglia

**DOI:** 10.1101/256347

**Authors:** David N. Lee, Apostolos P. Georgopoulos, Gert-Jan Pepping

**Affiliations:** Perception Movement Action Research Consortium, School of Philosophy, Psychology and Language Sciences and Moray House School of Education, University of Edinburgh, St Leonard’s Land, Holyrood Road, Edinburgh EH8 8AQ, UK; Brain Sciences Center, Veterans Affairs Medical Center, One Veterans Drive, Minneapolis, MN 55417, USA; Cognitive Sciences Center, University of Minnesota, Minneapolis, MN 55455, USA; Department of Neuroscience, University of Minnesota Medical School, Minneapolis, MN 55455, USA; Department of Neurology, University of Minnesota Medical School, Minneapolis, MN 55455, USA; Department of Psychiatry, University of Minnesota Medical School, Minneapolis, MN 55455, USA; School of Sport and Exercise Science, Australian Catholic University, 1100 Nudgee Road, Banyo, QLD 4014, Australia

## Abstract

Neural systems control purposeful movements both within an animal’s body (e.g., pumping blood) and in the environment (e.g., reaching). This is vital for all animals. The movement control functions of globus pallidus (GP), subthalamic nucleus (STN) and zona incerta (ZI) were analyzed in monkeys reaching for seen targets. Temporal profiles of their hand movements and the synchronized pattern of neuropower (rate of flow of electrochemical energy) through the basal ganglia were analyzed in terms of general tau theory of movement control (Lee et al., 2009), using the variable *rho* (=1/*tau*). The results suggest: (i) the neuroinformation for controlling movement is the relative-rate-of-change, *rho*, of neuropower in the nervous system; (ii) GP is involved in creating *prescriptive rhos* of neuropower to guide movements; (iii) STN is involved in registering perceptual *rhos* of neuropower to monitor the movement; (iv) ZI is involved in combining the prescriptive and perceptual *rhos* of neuropower to generate *performatory* rhos of neuropower to activate the muscles to produce the movement. Possible implications for Parkinson’s disease are discussed.

## Introduction

Purposeful movement, vital for the survival of any organism, is *controlled* by the organism’s nervous system. Three neurocontrol functions are involved: *perceptually* setting up and monitoring the movement, *prescribing* the movement, *performing* the movement. Because movements take time, these neurocontrol functions are all *prospective*, relating to the potential future course of the movement. In short, neurocontrol is based on prospective information.

Prospective control of movement involves guidance of body parts to goals across gaps so the parts move with appropriate momentum. For example, a seabird’s nervous system must prescribe a forceful movement of the bill to spear fish, but a much gentler movement of the bill when feeding the fish to its young. Likewise, when a cheetah is sprinting, its nervous system must prescribe forceful impact of the feet with the ground in order to achieve powerful thrusts, but when stalking, its nervous system must prescribe gentle approach of the feet to the ground so that it is not heard by the prey.

In this paper we shall report an experimental study of the function of the basal ganglia in rhesus monkeys in the prospective control of movement. The monkeys moved a handle to a seen goal, while the movement of the handle and the synchronous electrical activity in their basal ganglia were recorded. The study was based on an extended version of *general tau theory* of movement control (Lee et al. 2009), which was founded on the pioneering work of Gibson (1966) and Bernstein (1967). Principal tenets and implications of the extended theory are as follows:

i. *Movement-gaps*. Purposeful movement entails controlling the movement of body parts to goals across *movement-gaps*, from where one is to where one wants to be. Movement-gaps may be extrinsic or intrinsic to the organism, and across any physical dimension - e.g., distance gaps when reaching; angular gaps when looking; force gaps when gripping; intra-oral pressure gaps when suckling; pitch, loudness and timbre gaps when vocalizing.
ii. *ρ* (or *τ* = 1/*ρ*) is the primary informational variable used in controlling gaps. *ρ* of a gap equals the *relative rate of closing, or opening*, of the gap. (*τ* of a gap equals the time-to-closing, or time-from-opening of the gap at the current rate of closing, or opening.) Thus, *ρ* of gap *X* is

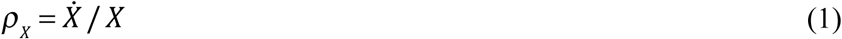

where the dot indicates the time derivative. *ρ*_*x*_ is directly perceptible through all known perceptual systems, in contrast with gap size, velocity or acceleration, which are not directly perceptible because they require scaling (Lee 1998).
iii. *ρ-coupling*. This enables the synchronous closing of two gaps, *Y* and *X*. It entails keeping

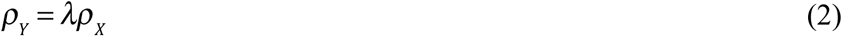

where *λ* is the coupling factor. *λ* determines the gentleness (if *λ* > 2) or forcefulness (if *λ* < 2) of the gap closure. For example, catching a ball gently or hitting it forcefully is achieved by keeping the rho of the hand to the catching place coupled to the rho of the ball to the catching place, with *λ* > 2 or *λ* < 2, respectively.
iv. *Stimulopower*. This is the rate of flow of energy from an external source to a receptor in an organism. The stimulopower may be power reflected from or emitted by a source. Stimulopower is what conveys information to an animal’s perceptual systems. In seeing, the stimulopower is electromagnetic; in hearing and echolocating, it is mechanical (vibrational); in touching, it is mechanical; in smelling, it is chemical; in heat-sensing, it is thermal; in electrolocating, it is electrical; in magnetosensing, it is magnetic.
v. *The elemental information in stimulopower*. When stimulopower is unattenuated by the medium through which it passes it is proportional to the power emitted by the source (*ρ*_*source*_) divided by the square of the size of the gap (*r*) between of the source and the receptor. Thus

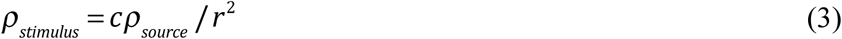 This means that gap-size, *r*, is *not* specified by *ρ*_*stimulus*_ unless *ρ*_*source*_ and the constant of proportionality, *c*, are known, which we may assume is not normally the case. Therefore, no time derivative of gap-size (velocity, acceleration, jerk, etc) is specified either. However, when *ρ*_*source*_ is constant during the registering of stimulopower then, from Eqns. (1) and (3), y

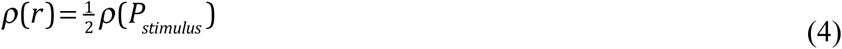
vi. *Derivative information about relative gap size.* Although the rho of stimulopower incident on a receptor does not provide information about the size of the gap to the source of the stimulopower, the rhos of stimulopower from two or more directions falling on an array of receptors, as on a retina, can provide information about relative gap sizes (Lee et al. 2009, Fig.2).
vii. *Pain*. When the stimulopower exceeds the pain threshold, *P*_*stimulus*_ is registered not by sensoriceptors but by nociceptors, which pick up information for controlling the closing or opening of a potentially harmful motion-gap, *r*. The equation is

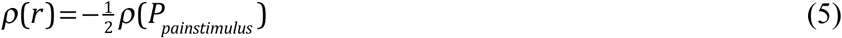
viii. *Stimulopower to neuropower*. At a sensori-receptor, or nociceptor, stimulopower is transduced into *neuropower* (rate of flow of electrochemical energy through the nervous system). The transduction follows a mathematical power law. This means that the *ρ* of neuropower is proportional to the *ρ* of stimulopower (Lee et al. 2009). If the power exponent is unity, *ρ* of neural power equals *ρ* of stimulopower. In the periphery, the *ρ* of neuropower appears as the *ρ* of graded electrical potentials in nerves. More centrally, in neural axons, the *ρ* of neuropower appears as trains of action potentials of about uniform energy (Kandel *et al.* 2000). These are produced by sodium/potassium pumps injecting bursts of ionic energy at nodes of Ranvier to speed transmission of the neuropower. Thus, the *ρ* of neuropower in axons appears as the *ρ* of the rate of flow of electrical spikes. When an axon synapses on a dendrite of another cell, that cell releases a neurotransmitter and the electrical neuropower in the axon is power law transduced into chemical neuropower. Thus the *ρ* of the chemical neuropower is proportional to the *ρ* of the electrical neuropower. Post-synaptically, the chemical neuropower is power law transduced into ionic neuropower in dendrites (recorded as graded synaptic potentials), again following a power law transduction process. Finally, the rhos of the electrical neuropower in the dendrites is spatiotemporarily aggregated in the cell body, resulting in an aggregate *ρ* of electrical neuropower flowing in the cell’s axon, in the form of a train of action potentials. And so the process continues.
ix. *Prescriptive neuropower*. To achieve purposeful movement, an organism’s nervous system must *prescribe* the dynamic form of the movement to be performed and continuously modify the prescription on the basis of perceptual information about how the movement is actually proceeding. To do this, the nervous system prescribes the *ρ* of a movement-gap by generating a special *ρ*, *ρ*_*G*_, and coupling the *ρ* of the gap to be controlled, *ρ*_*Y*_, onto *ρ*_*G*_. Thus,

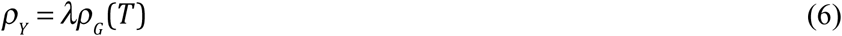

where *T* is the duration of the gap-closing movement (from zero), and *λ* specifies the velocity profile of the movement (Fig. 1). The equation is

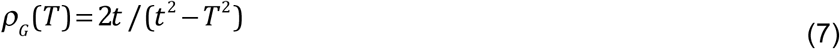

where time, *t*, runs from zero to *T*.
x. *ρ*_*G*_ is rooted in gravity, an ecological invariant for all animals. *ρ*_*G*_ is the *ρ* of a movement-gap that changes from rest at constant acceleration, like something dropping under gravity. This means that animals are often stimulated by *ρ*_*G*_. For example, when an animal is running, during every stride it alternates between being in free-fall under gravity and being supported by the ground. As it passes from free-fall to support, mobile masses within body cavities, notably the otoliths in the saccules of vertebrate vestibular systems, accelerate at a constant rate downward under gravity. Thus, the motion of the mass (otolith) relative to the cavity (saccule) follows the *ρ*_*G*_ function. Therefore, if there are sensors to detect the motion of the mass, as there are in the saccule, the animal will constantly receive *ρ*_*G*_ stimulation as it runs along.
xi. *Evidence for ρ_G_-control*. The process of using *ρ*_*G*_ to prescribe movement-gaps is called *ρ*_*G*_-*control*. Experiments indicate *ρ*_*G*_-control of movement-gaps by (i) human adult*s* when looking, reaching, swaying, walking, sprinting, golfing, playing music, singing, speaking (summarized in Lee et al. 2009); (ii) Parkinson’s patients when controlling sway and being helped by listening to and then remembering *tauG* - controlled sounds (Schogler et al 2017); (iii) human neonates when controlling intraoral pressure when suckling (Craig & Lee 1999); (iv) birds and bats when steering and landing (Lee et al. 2009). There is also evidence for *ρ* in the brains of locusts (Rind & Simmons 1999), pigeons (Sun and Frost 1998), monkeys (Merchant et al. 2004; Merchant & Georgopoulos, 2006) and humans (Field & Wann, 2005; Tan et al., 2009; van der Weel *et al.* 2009). We investigated the neuroprocesses underpinning *ρ*_*G*_-control by analyzing single unit recordings from external and internal globus pallidus (GPe, GPi), subthalamic nucleus (STN) and zona inserta (ZI), when monkeys were moving their hand to a goal along a straight track. These basal ganglia are thought to be involved in sensorimotor control (Mitrofanis 2005; Fasano et al. 2015; Takamitsu & Yamomoto 2015)). There is also evidence of temporal coherence in basal ganglia during voluntary movement (Talakoub et al. 2016). However, how the temporal pattern of neuropower in basal ganglia relates to the temporal pattern of voluntary movement has not been studied. This was our aim.

**Fig. 1.**
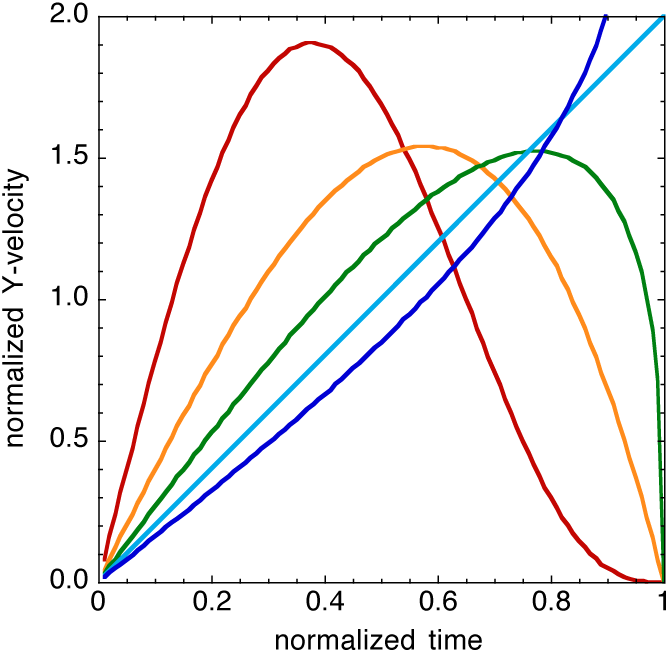
Velocity of closure of a movement-gap, *Y*, that is *ρ*_*G*_-guided following the equation *ρ*_Y_ = *λ ρ*_*G*_ (*T*). Effect of the value of coupling factor, λ, on the velocity of closure of *Y*: *λ*= 4 (red); *λ*= 2 (orange); 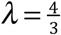 (green); *λ*= 1 (light blue); 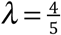 (blue), Amplitude and duration of the movement have each been normalized to 1 for illustration.

## RESULTS

### Relation between neuropower and hand movement

On each trial, the neuropower in GPi, GPe, STN and ZI was measured as spike-rate, the rate of flow of action potentials. The neuropower was time locked onto the start of the hand movement and time-normalized into bins of duration 0.05 times the duration of the hand movement on that trial; i.e., into bins of duration 0.05 movement-time units (mtus). The mean and standard error of the time-normalized neuropower time series were then computed across all the trials in each ganglia under study. The black lines in Figure 2A show the means and standard errors of the time-normalized neuropowers. The coloured lines show these neuropowers smoothed with a Gaussian filter, sigma 0.1 mtu.

**Fig. 2.**
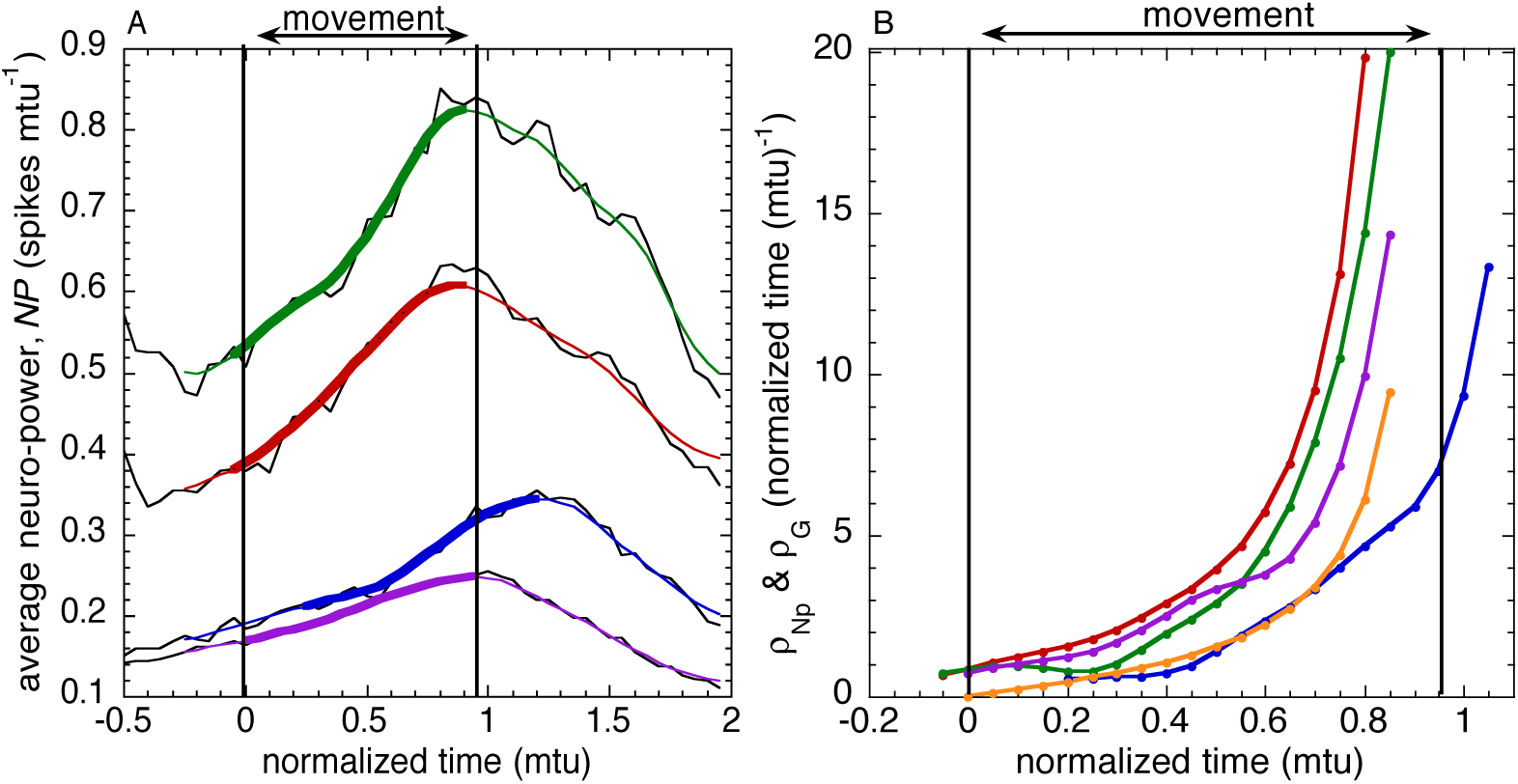
(A) Neuropower profiles during hand movement. The neuropower profiles were measured as spike-rate and were time-normalized with respect to hand movement time and time-locked to it. Black lines: means of unsmoothed neuropower. Coloured lines: Gaussian (sigma 0.1 mtu) smoothed values of GPe (red), GPi (green), STN (blue), ZI (purple). Thicker coloured lines: the sections of the neuropower profiles that were of the same duration as the hand movement (1 mtu), and the *ρ* s of which were the most highly correlated with *ρ*_*Mov*_, the *ρ* of the hand movement. (B) The *ρ* s of the neuropower sections in (A), together with *ρ*_*G*_ (orange line).

Even though the means and standard errors of the time-normalized neuropowers (black lines in Fig. 2A) were computed across many different neurons (214 in GPe, 86 in GPi, 33 in STN, 68 in ZI) and in three monkeys (Methods), the standard errors (se) were remarkably small. As a fraction of the maximum value of neuropower, the mean ± se of the standard errors of mean neuropower were: 0.00157 ± 0.00001 (GPe), 0.00108 ± 0.00001 (GPi), 0.00123 ± 0.00003 (STN), 0.00121 ± 0.00001 (ZI). These small standard errors (possibly related to the monkeys’ hand movements on the task being well practiced) strongly suggest that the average duration-independent neuropower (black lines in Fig. 2A) measures a basic time-invariant of the hand movement, and thereby provides a measure of the average neuroactivity (across neurons in a tract) underpinning the hand movement.

The sections of the mean time-normalized neuropower profiles (Fig.2A) that were most strongly *ρ*-coupled to the hand movement were determined by computing, for each time-normalized neuropower profile, the *ρ* s of the neuropower sections of duration 1 mtu (movement time unit) that ended at each point in the neuropower profile. The *ρ* of each of these sections of neuropower, *ρ*_*NP*_, was then linearly regressed on *ρ*_*Mov*_, the rho of the movement up to the goal position, and the section of the neuropower profile that yielded the highest *r*^*2*^ was taken as corresponding to the hand movement. These sections of the neuropower profile were found to end at the peak mean time-normalized neuropower in each ganglia studied. In GPe and GPi, the peak occurred 0.05mtu (on average about 25ms) before the movement ended. In ZI it occurred just as the movement ended. In STN it occurred 0.2mtu (on average about 100ms) *after* the movement ended.

### Coupling *ρ*_*NP*(*GPe*)_ onto *ρ*_*G*_

The degree to which the *ρ* of neuropower (*ρ*_*NP*_) was rho-coupled onto *ρ*_*G*_ was measured by plotting *ρ*_*NP*_ against *ρ*_*G*_ for each of the four ganglia, and calculating the linear regressions forced to pass through the origin (since *proportionate* coupling was predicted). The regression lines are shown in Fig. 3A. The regression coefficients are given in Table 1. The *R*^*2*^ values, the coefficients of determination, measure the strengths of the rho-couplings, i.e., the proportion of variance in *ρ*_*NP*_ accounted for by the rho-couplings. The strength of the rho-coupling of *ρ*_*NP*(*GP*_*e*_)_ onto *ρ*_*G*_ was very high (*R*_2_ = 0.996, compared with a maximum possible value of 1.000). This strongly suggests that GPe was involved in creating the prospective-guiding function, *λρ*_*G*_. The slopes of the regressions (Table 1) measure the *λ* coupling factors. For *ρ*_*NP(GPe)*_ on *ρ*_*G*_, *λ* was 2.13, which indicates that the neuropower in GPe approached its peak value gently, stopping when it got there (c.f. Fig.1).

**Table 1.** Degrees of rho-coupling (r^2^ of linear regressions through origin). In parentheses, rho-coupling factors (the regression slopes). Red, r^2^≥0.990; purple, r^2^≥0.980. Rows, dependent variables; columns, independent variables (e.g., *ρ*_*NP(GPE*)_ = 2.13*ρ*_*G*_

|  |  |  |  |  |  |  |
| --- | --- | --- | --- | --- | --- | --- |
| $\rho_{NP(GPe)}$ | 0.996<br>(2.13) | | 0.990<br>(1.312) | 0.988<br>(1.316) | 0.969<br>(1.343) | 0.965<br>(0.278) |
| $\rho_{NP(GPi)}$ | 0.986<br>(1.615) | 0.989<br>(0.758) | | 0.974<br>(0.996) | 0.963<br>(1.019) | 0.954<br>(0.219) |
| $\rho_{NP(ZI)}$ | 0.973<br>(1.604) | 0.987<br>(0.755) | 0.970<br>(0.988) | | 0.980<br>(1.020) | 0.982<br>(0.214) |
| $\rho_{NP(STN)}$ | 0.960<br>(1.559) | 0.965<br>(0.732) | 0.956<br>(0.961) | 0.980<br>(0.970) | | 0.989<br>(0.212) |
| $\rho_{Mov}$ | 0.984<br>(7.358) | 0.969<br>(3.546) | 0.948<br>(4.453) | 0.975<br>(4.609) | 0.986<br>(4.687) | |
| | $\rho_G$ | $\rho_{NP(GPe)}$ | $\rho_{NP(GPi)}$ | $\rho_{NP(ZI)}$ | $\rho_{NP(STN)}$ | $\rho_{Mov}$ |

**Fig. 3.**
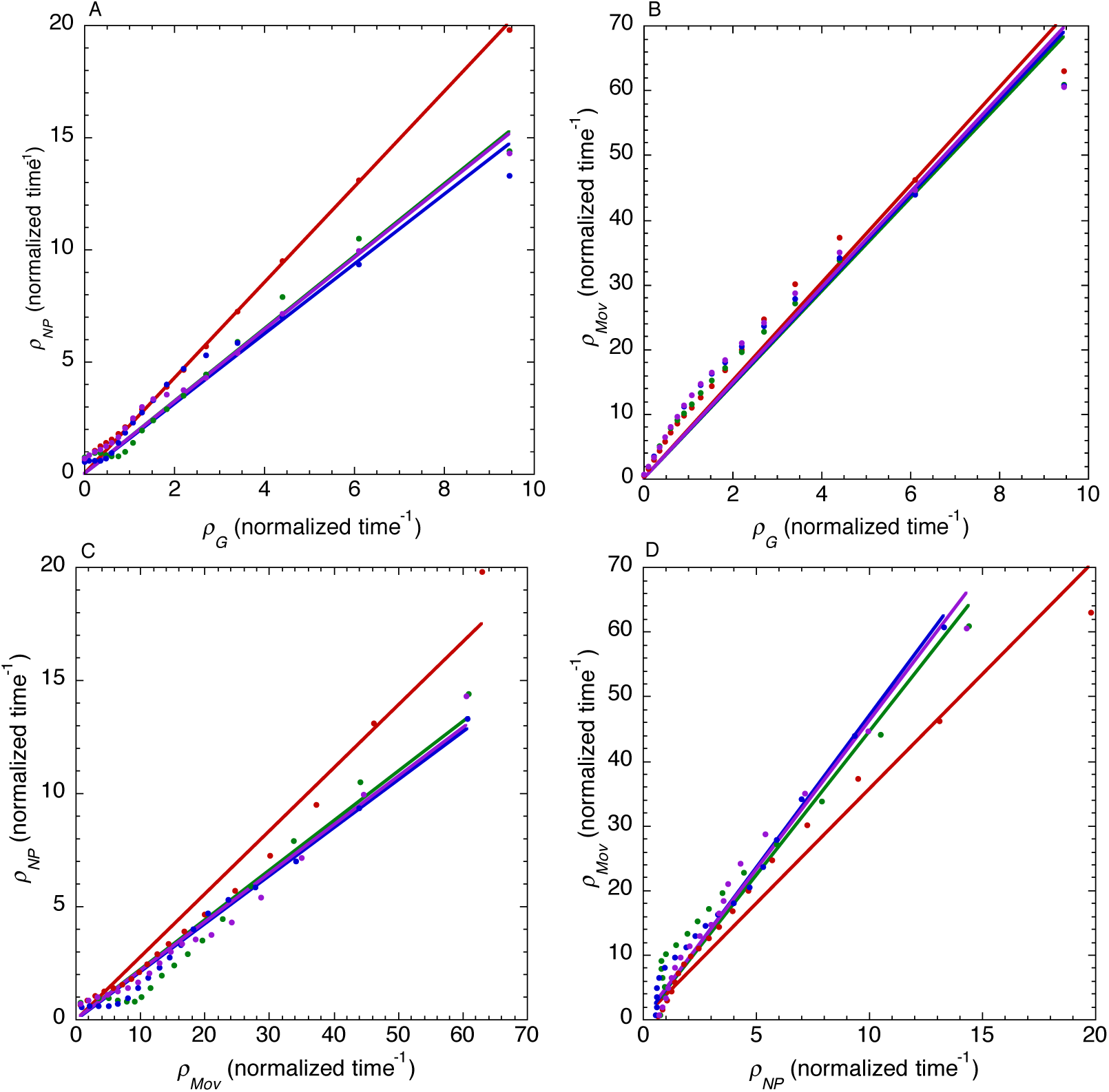
Rho-coupling regressions through origin. (A) Rho of neuropower (*ρ*_*NP*_) on hypothesized prospective rho (*ρ*_*G*_). (B) Rho of hand movement *ρ*_*Mov*_, on *ρ*_*G*_. (C) *ρ*_*NP*_ on *ρ*_*Mov*_. (D) *ρ*_*Mov*_ on *ρ*_*NP*_.

**Fig. 4.**
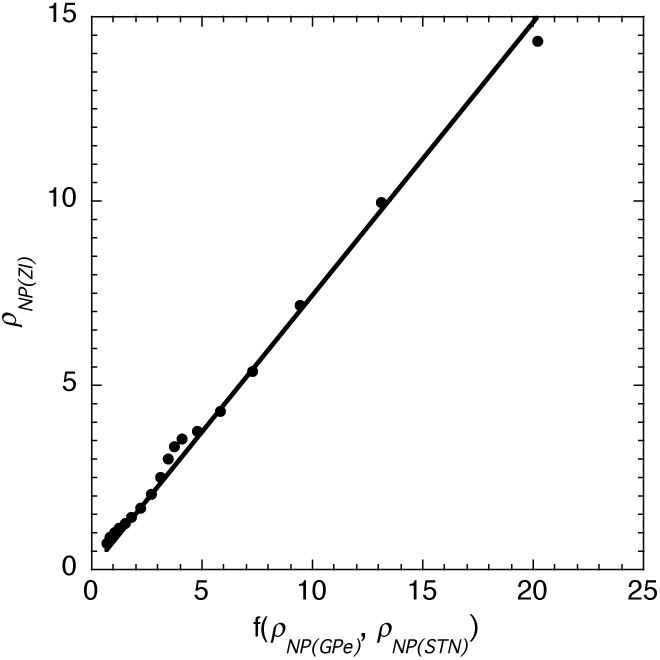
*ρ* of neuropower in ZI. *ρ*_*NP(ZI*)_, is plotted against the *ρ* input to ZI, *ƒ*(*ρ*_*NP*(*GPe*)_,*ρ*_*NP*(*STN*)_; see equations (4) and (5). The linear regression line was forced to pass through the origin to test predicted *proportional ρ*-coupling. The regression equation was *ρ*_*NP*(*ZI*)_ =0.740 *ƒ*(*ρ*_*NP*(*GPE*)_,*ρ*_*NP*(*STN*)_),*R*^*2*^=0.993.

### Coupling *ρ*_*Mov*_ onto *ρ*_*G*_

The degree to which the *ρ* of the movement (*ρ*_*Mov*_) was rho-coupled onto *ρ*_*G*_ was measured by plotting the average *ρ*_*Mov*_ across the four ganglia studied and calculating the linear regressions through the origin (since *proportionate* coupling was predicted). The regressions are shown in Fig. 3B. The regression coefficients are given in Table 1. The *R*^*2*^ for the coupling of *ρ*_*Mov*_ onto *ρ*_*G*_ was 0.984. This indicates that *ρ*_*Mov*_ followed *ρ*_*G*_ quite closely. The coupling factor was 7.358, indicating that the hand approached the target very gently, slowing down quite early to stop at it (c.f. Fig. 1).

### Coupling *ρ*_*NP*_ onto *ρ*_*Mov*_

The degree of coupling of *ρ*_*NP*_ onto *ρ*_*Mov*_ was measured by plotting *ρ*_*NP*_ against *ρ*_*Mov*_ for each ganglion and calculating the linear regressions through the origin (since proportionate coupling was predicted). The regressions are shown in Fig. 3C. The regression coefficients are given in Table 1. The neuropower section in STN that corresponded to the hand movement started 0.2 mtu *after* the hand movement started. The highest *R*^*2*^ (0.989) was for STN. Taken together these findings suggest that STN was involved in the perceptual monitoring of *ρ*_*Mov*_, after a perceptual delay of 0.2 mtu (about 100 ms on average).

### Coupling *ρ*_*Mov*_ onto *ρ*_*NP*_

The degree of coupling of *ρ*_*Mov*_ onto *ρ*_*NP*_ was measured first by plotting *ρ*_*Mov*_ against *ρ*_*NP*_ for each ganglion studied and calculating the linear regressions through the origin. The regressions are shown in Fig. 3D. The regression coefficients are given in Table 1. The two highest *R*^2^ were for *ρ*_*NP(STN*)_ (0.986) and *ρ*_*G*_ (0.984). This suggested to us that *ρ*_*NP(STN*)_ and *ρ*_*G*_ might have been jointly influencing *ρ*_*Mov*_. In particular, that ZI might receive movement-adjusting *ρ* information from STN and GPe (since *ρ*_*NP(GPE*)_ was very strongly coupled to *ρ*_*G*_; *r*^*2*^ = 0.996) and transmit this information to the muscles, and thence to the hand movement. To investigate this possibility we defined 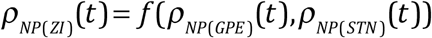, and then sought to determine the function *f* from the experimental data. Taking into account the observed perceptual delay of 0.2 mtu, when STN could not have registered the hand movement, we hypothesized that, for t = 0 to 0.2 mtu,

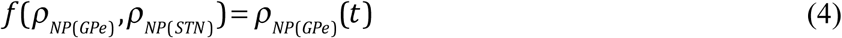

and for t = 0.2 to 1.0 mtu,

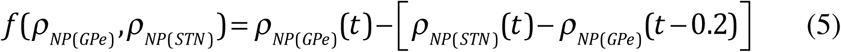

where 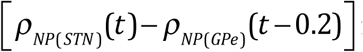 is the deviation of the movement as perceived from its prospective rho course. To test the hypothesis we computed the linear regression (through the origin) of *ρ*_*NP(ZI*)_ on 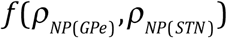 for t = 0 to 1.0 mtu. The regression yielded an *R*^*2*^ of 0.993 and a coupling factor of 0.740. Thus, the hypothesis was strongly supported.

## DISCUSSION

We have argued that animals prospectively control their movements by using, as information in the neurosystem, the relative-rate-of-change, *ρ*, of neuropower (i.e., the rate of flow of electrochemical energy through neurons). A principal argument that *ρ* is the fundamental informational variable for controlling movement is that the *ρ* of the distance gap that needs to be closed to achieve a controlled movement is directly perceptible, whereas the distance itself, or any of its time derivatives, are not directly perceptible.

The results of our analysis of the electrical activity in basal ganglia GPe, GPi, STN and ZI of monkeys reaching to seen targets is consistent with the idea that GPe is implicated in *prospectively guiding* the *ρ* of a movement using the prescriptive neuropower function, *ρ*_*G*_; that STN is implicated in the *perceptual monitoring* of the movement; and that ZI is implicated in integrating (Sherrington 1961, Branco et al 2010, 2012) the *ρ* of *prospective* neuropower from GPe and the *ρ* of *perceptual* neuropower from STN to create the *ρ* of *enacting* neuropower at ZI. The *ρ* of the *enacting* neuropower both informationally and physically (after amplification, e.g., with ATP) powers the *ρ* of *muscular-power* (the rate of flow of energy into the muscles) and this powers the *ρ* of the movement. Thus the *ρ* of power has come full circle.

Perhaps it is not too surprising that the basal ganglia should be implicated in the three fundamental neurofunctions controlling movement - prospecting, perceiving and performing. After all, basal ganglia are phylogenically ancient in vertebrates - including those, like lamprey, who lack cerebral cortices (Grillner 2003) – and all vertebrates control their movements purposefully in order to live.

### Parkinson’s Disease

The motor symptoms of Parkinson’s Disease - tremor, rigidity, bradykinesia, freezing, and dysarthria - are generally considered to involve malfunction in the basal ganglia (Moustafa, et al., 2016). However, the electrophysiological functions in the basal ganglia that are affected are unknown (Ellens & Leventhal, 2013). The present study could cast some light on the issue. Many movements are affected in Parkinson’s disease, but some are relatively unaffected - the so-called paradoxical movements. For example, catching a moving ball can be easier than reaching for a stationary one, and walking downstairs can be easier than walking across a featureless floor. The difference in ease of performance could be related to the type of information being used. The information for catching a moving ball or walking downstairs is largely perceptual, from the optic flow field at the eye; whereas, when reaching for a stationary ball or walking across a featureless floor, movement control is more reliant on prospectively-guiding information created in the neurosystem. Since our results indicate that GPe is strongly implicated in generating prospectively-guiding *ρ*_*G*_ information, it is possible that movement disorders in Parkinson’s –tremor, rigidity, bradykinesia, freezing, and dysarthria - which all involve poor prospective coordination of muscles – may be due, in part at least, to GPe dysfunction.

### Where next?

A principal function of any neurosystem is controlling bodily movements. If, as our results suggest, the common informational currency in basal ganglia for controlling movements is the *ρ* of neuropower, then it is likely that the same informational currency is used throughout the neurosystem when movements are being controlled (otherwise a Tower of Babel situation would prevail). This idea could be tested in humans, for example, by using high temporal resolution MEG (Tan et al., 2009). If verified, rho theory might then be used to analyze normal function in the neurosystem, and also reveal regions of the brain where there is dysfunction in the transmission of information for guiding movement.

Rho theory might also be useful in investigating how other organisms control their movements. For example, Delafield-Butt *et al.* (2011) have obtained motion-capture evidence that unicellular paramecia guide their movement to an electric pole using the *ρ*_*D*_ function as a guide; Strausfeld and Hirth (2015) have suggested that the central complex in an insect’s brain is homologous with the basal ganglia in animals, and so might have similar control functions; and Darwin and Darwin (1880) and Masi *et al.* 2009 have suggested that the transition zone in the roots of plants is implicated in neurocontrol of movement. It would be of value to investigate to what extent the *ρ* of neuropower is used generally by organisms in controlling their movements.

## Methods

We analyzed neural and movement data from three rhesus monkeys in four experiments. In each experiment there were two horizontal rows of 128 LEDS, 32 cm long, one above the other. The monkeys were trained to move a handle to the left or right to line up an LED on the lower row with a target LED on the upper row. The handle was constrained to run along a track 32 cm long. The position of the handle was recorded every 10 ms. The movements averaged 525 ms and 248 mm. Neural electrical activity was recorded extra-cellularly with microelectrodes on separate occasions from five hemispheres of three monkeys: from 214 cells in the arm area of globus palidus external (GPe), 86 cells from globus pallidus internal (GPi), 33 cells in the subthalamic nucleus (STN) and from 75 cells in zona incerta (ZI). Details of the procedures for the experiments are given in DeLong et al. (1985) and Crutcher et al. (1980)

### Data analysis

The data for GPe, GPi, STN and ZI were first assembled into *unit records*, which comprised the hand position data and neural data recorded on a single reach-to-target trial. Using the index *i* to refer to a unit record, and the index *j* to refer to a 10 ms time sample, a unit record comprised (i) a hand position time series, *x*_*i,j*_, where *x* is the coordinate of the handle, and (ii) a neural spike-density time series, *n*_*i,j*_, where *n*_*i,j*_ is the number of neural spikes in the *jth* 10 ms time bin of the *ith* unit record. Then, for each unit record (i) to minimize noise in the data, the *x*_*i,j*_. time series of the hand were smoothed, with a Gaussian filter with time constant sigma of 30 ms, and a cutoff frequency of 6 Hz, yielding the smoothed time series *X*_*i,j*_; (ii) the *X*_*ij*_ time series were numerically differentiated with respect to time to yield the time series 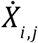; (iii) the movement time, *MT*_*i*_, for the *ith* unit record was calculated as the difference between the ‘hand-start’ and ‘hand-end’ times, defined respectively as the first sample time after the speed of the hand rose above and the last sample time before it fell below 5% of the peak speed on that recording. *MT*_*i*_ averaged 525ms; (iv) each hand/target action-gap time series, *M*_*i,j*_, was computed as the distance between the position of the handle at each sample time, *j*, and its position at the end of the movement.

A unit record was accepted for further analysis providing the neuron became ‘active’ between the stimulus light going on and the hand starting to move; i.e., providing there were five or more consecutive 10 ms bins in the time series which were of value greater than three standard deviations above the mean value of *n*_*i,j*_ during the 500 ms preceding the stimulus light. This criterion was applied to exclude normalized unit records where there was no evidence of the neural spike-rate being related to the hand movement. In the records satisfying the criterion (i) the members of the hand/target gap time series, *M*_*i,j*_, were numerically differentiated with respect to time to yield the time series 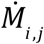; (ii) the time series, *ρ*_*i,j*_ of the hand/target gap, was calculated using the formula 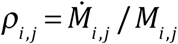 (iii) the time series 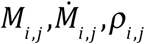 were time-normalized to yield the time series 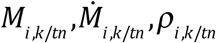. Time-normalization entailed apportioning the data in each time series into equal time bins of width *MT*_*i*_/20 s, or 0.05 mtu (movement time units), so that for the movement of the hand between the start time and the end time the index, *k*, ran from 0 to 19, and normalized time ran from 0 mtu to 0.95 mtu. For the *n*_*i,k/tn*_ time series the index *k* ran from −40 through 0 (when the hand movement started) to +39, and the corresponding normalized time ran from −2.00 mtu through 0 (when the hand movement started) to +1.95 mtu. Normalizing time in this way meant that the normalized duration of the hand movement remained the same across the normalized records, enabling the average time-normalized spike-density function, which was assumed to be coupled in *relative* time to the hand movement, to be measured as the mean time series, 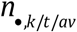 of the *n* time series, for *k =* −40, −39, ….39, and normalized time running from −2.00 mtu to 1.95 mtu. The values of 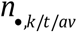 were computed separately for the GPe, GPi, STN and ZI time-normalized records. The time series 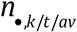 were also averaged separately across the records, yielding the mean time series, 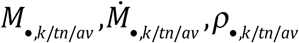 for *k* = 0, 2, ….19, and normalized time running from 0 mtu to 0.95 mtu.

To enable statistical comparisons to be made, the same analysis was performed on 1000 samples, drawn with replacement from each of the normalized records. This resulted, for each brain area, in 1000 sets of average time series 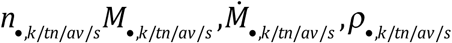, for s=1, 2, … 1000.

### Summary: time series used in the analysis

In summary, the time series used in analyzing the results were 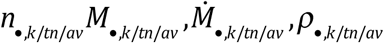. In the main text and figures these average normalized-time time series are, for conciseness, designated respectively as *n* (the average time-normalized spike density function, which is taken as the measure of average *neuropower*), *M* (the average time-normalized action-gap), 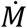 (the time derivative of the average time-normalized action-gap), and *ρ*_*M*_ (the *ρ* of the average time-normalized action-gap). These time series are graphed in the figures for normalized time extending from −2 mtu to +2 mtu in steps of 0.05 mtu, which corresponds to *k* = 1, 2,………80.

## Acknowledgements

The research was supported by grants from American Legion Brain Sciences Chair, BBSRC, EU NEST-ADVENTURE, Leverhulme Trust, US Public Health Service Grant PSMH48185, US Department of Veterans Affairs.

